# Transcriptome analysis in the silkworm *Bombyx mori* overexpressing piRNA-resistant *Masculinizer* gene

**DOI:** 10.1101/2022.05.13.491740

**Authors:** Kenta Tomihara, Susumu Katsuma, Takashi Kiuchi

## Abstract

Dosage compensation is a process that produces a similar expression of sex-linked and autosomal genes. In the silkworm *Bombyx mori* with a WZ sex-determination system, the expression from the single Z in WZ females matches that of ZZ males due to the suppression of Z-linked genes in males. A primary maleness determinant gene, *Masculinizer* (*Masc*), is also required for dosage compensation. In females, P-element induced wimpy testis (PIWI) is complexed with the W chromosome-derived female-specific *Feminizer* (*Fem*) PIWI-interacting RNA (piRNA) and cleaves *Masc* mRNA. When *Fem* piRNA-resistant *Masc* cDNA (*Masc-R*) is overexpressed in both sexes, only female larvae are dead during the larval stage. In this study, transcriptome analysis was performed in neonate larvae to examine the effects of *Masc-R* overexpression on a global gene expression profile. Z-linked genes were globally repressed in *Masc-R*-overexpressing females due to force-driven dosage compensation. In contrast, *Masc-R* overexpression had little effect on the expression of Z-linked genes and the male-specific isoform of *B. mori insulin-like growth factor II mRNA-binding protein* in males, indicating that excessive *Masc* expression strengthens neither dosage compensation nor maleness in males. Fourteen genes were differentially expressed between *Masc-R*-overexpressing and control neonate larvae in both sexes, suggesting Masc functions other than dosage compensation and masculinization.

**Highlights:**

- Transcriptome analysis was performed in *Masc-R*-overexpressing neonate *Bombyx mori* larvae.
- Z-linked genes were globally suppressed in *Masc-R*-overexpressing females.
- *Masc-R* overexpression had little effect on *BmImp^M^* expression in males.
- Several genes may be controlled by *Masc-R* regardless of sex.

## Introduction

Genes that function at the uppermost stream of the sex-determination cascade also regulate dosage compensation, which has the effect of equalizing expression of sex-linked and autosomal genes. For example, in the fruit fly *Drosophila melanogaster* that uses an XY sex-determination system, a dosage compensation complex activates the transcription of X-linked genes in XY males. The master femaleness determinant protein, sex-specific lethal (SXL), targets *male-specific lethal 2* (*msl2*) mRNA and represses its translation in XX females [1–3]. Because *msl2* encodes a subunit of the dosage compensation complex, dosage compensation occurs only in the absence of SXL (i.e., XY males). The nematode *Caenorhabditis elegans* uses an XO sex-determination system, and genes on both X chromosomes are equally suppressed by a dosage compensation complex in XX hermaphrodites. In *C. elegans*, dosage compensation occurs only in the hermaphrodites because the male determinant protein XO lethal (XOL-1) suppresses *sex determination and dosage compensation protein* (*sdc-2*) expression in XO males, which encodes a subunit of the dosage compensation complex [4,5]. In contrast, in eutherian mammals that use an XY sex-determination system, sex-determination and dosage compensation are achieved by two independent pathways [6].

The silkworm *Bombyx mori*, a model species of lepidopteran insects, uses a WZ sex-determination system [7]. The expression from the single Z in WZ females matches that of ZZ males because the expression of Z-linked genes is suppressed in males [8,9]. Kiuchi et al. [10] reported that a Z-linked gene *Masculinizer* (*Masc*) is essential for maleness and dosage compensation. In WZ females, *Masc* mRNA is cleaved by silkworm P-element induced wimpy testis (PIWI)-like protein (Siwi) complexed with W-linked *Feminizer* (*Fem*) PIWI-interacting RNA (piRNA) [10]. In *Masc* knocked down males, most Z-chromosome-derived transcripts are transcriptionally upregulated [8,10], showing that Masc reduces Z-linked gene expression at the embryonic stage in ZZ males.

Previously, transfection of *Fem* piRNA-resistant *Masc* (*Masc-R*) cDNA resulted in growth inhibition of *B. mori* ovary-derived cultured cells [10,11]. In *Masc-R* cDNA-transfected cultured cells, the expression levels of autosomal genes were comparable to those in control cells, whereas the expression levels of Z-linked genes slightly decreased [11]. Sakai et al. [12] reported that *B. mori* females overexpressing *Masc-R* under the control of the *B. mori cytoplasmic actin 3* (*BmA3*) promoter were dead during the larval stage. However, the effects of *Masc-R* overexpression on dosage compensation in living females were not investigated. In addition, although *Masc-R*-overexpressing males did not show any abnormalities in growth, development, and sexual dimorphisms [12], whether *Masc-R* overexpression strengthens the dosage compensation and “maleness” in males has not been clarified yet.

In this study, transcriptome analysis was performed in *Masc-R*-overexpressing neonate larvae to examine the effects of *Masc-R* overexpression on a global gene expression profile. Z-linked genes were repressed in *Masc-R*-overexpressing females. In contrast, the expression of Z-linked genes and a masculinizing marker, the male-specific isoform of *B. mori insulin-like growth factor II mRNA-binding protein* (*BmImp^M^*) [13] in *Masc-R*-overexpressing males, was comparable to that in control males. Additionally, genes directly or indirectly regulated by Masc in both sexes were identified.

## Materials and Methods

### Insects

The ubiquitous *GAL4*-expressing strain BmA3-GAL4 [14] and the *UAS-Masc-R* carrying strain Sumi13-3 [12] were gifts from Drs. Megumi Sumitani and Hideki Sezutsu (National Agriculture and Food Research Organization). A BmA3-GAL4 female was crossed to a Sumi13-3 male, and F1 neonate larvae expressing *Masc-R* transgene were obtained by selecting double fluorescent-positive larvae that expressed both DsRed (*GAL4* marker) and enhanced green fluorescent protein (*UAS* marker) in their eyes. All larvae were reared on mulberry leaves or an artificial diet [SilkMate PS (Nosan)] under 12 h light/12 h darkness conditions at 25°C.

### RNA extraction, molecular sexing, and reverse transcription-quantitative polymerase chain reaction (RT-qPCR)

*Masc-R*-overexpressing neonate larvae (*A3-GAL4/+; UAS-Masc-R/+*) and control neonate larvae (+/+; +/+) were sampled from the same batch. Both DNA and RNA were extracted from a whole body of neonate larva with TRIzol (Thermo Fisher Scientific) according to the previous study with some modifications [15]. Genomic PCR for molecular sexing was performed using KOD One (Toyobo) with the primer sets listed in Table S1. The number of individuals used in further experiments is shown in Table S2. Reverse transcription was performed using the oligo(dT) primer and avian myeloblastosis virus reverse transcriptase contained in the TaKaRa RNA PCR kit (TaKaRa). RT-qPCR was performed using the Kapa SYBR FAST qPCR kit (Kapa Biosystems) and StepOnePlus™ Real-time PCR System (Applied Biosystems) with the primer sets listed in Table S1.

### RNA sequencing (RNA-seq) for differential expression (DE) analysis in *Masc-R*-overexpressing larvae

Larval total RNA was pooled together for each experimental group, and libraries for RNA-seq were generated. The libraries were sequenced by the NovaSeq 6000 Sequencing System (Illumina) with 100 bp paired-end reads. Library preparation and sequencing were performed by Novogene (Beijing, China). Adapter sequences and low-quality reads were removed with Trim Galore! (https://www.bioinformatics.babraham.ac.uk/projects/trim_galore/). The trimmed reads were mapped to the *B. mori* reference genome [16] with STAR [17], and the expression levels for each gene were estimated by RSEM [18]. DE analysis was performed by edgeR using read counts normalized with using the trimmed mean of M-values (TMM) method [19]. Genes with counts per million (CPM) values of <1 in both libraries were filtered out in the DE analysis. Using differentially expressed genes [DEG; false discovery rate (FDR) < 0.05], Gene Ontology (GO) enrichment analysis was performed by the GOseq R package [20]. GO terms corresponding to each gene model were obtained from KAIKObase (https://kaikobase.dna.affrc.go.jp) [21].

## Results

This study obtained neonate larvae that possess the *UAS-Masc-R* transgene [12] and the *GAL4* gene under the control of the *BmA3* promoter [14]. RT-qPCR showed that the expression level of *Masc* was ~75-fold higher in *Masc-R*-overexpressing females than in control females and ~50-fold higher in *Masc-R*-overexpressing males than in control males (Fig. 1A). *BmImp^M^* expression [13] was induced in *Masc-R-*overexpressing females (Fig. 1B), indicating that the *Masc-R* mRNA level in the *Masc-R*-overexpressing strain is sufficient for inducing the masculinizing cascade in females [12]. In contrast, the expression level of *BmImp^M^* in *Masc-R*-overexpressing males is comparable to that of control males (Fig. 1B), showing that an excessive amount of Masc protein does not enhance masculinization in males.

**Fig. 1.**
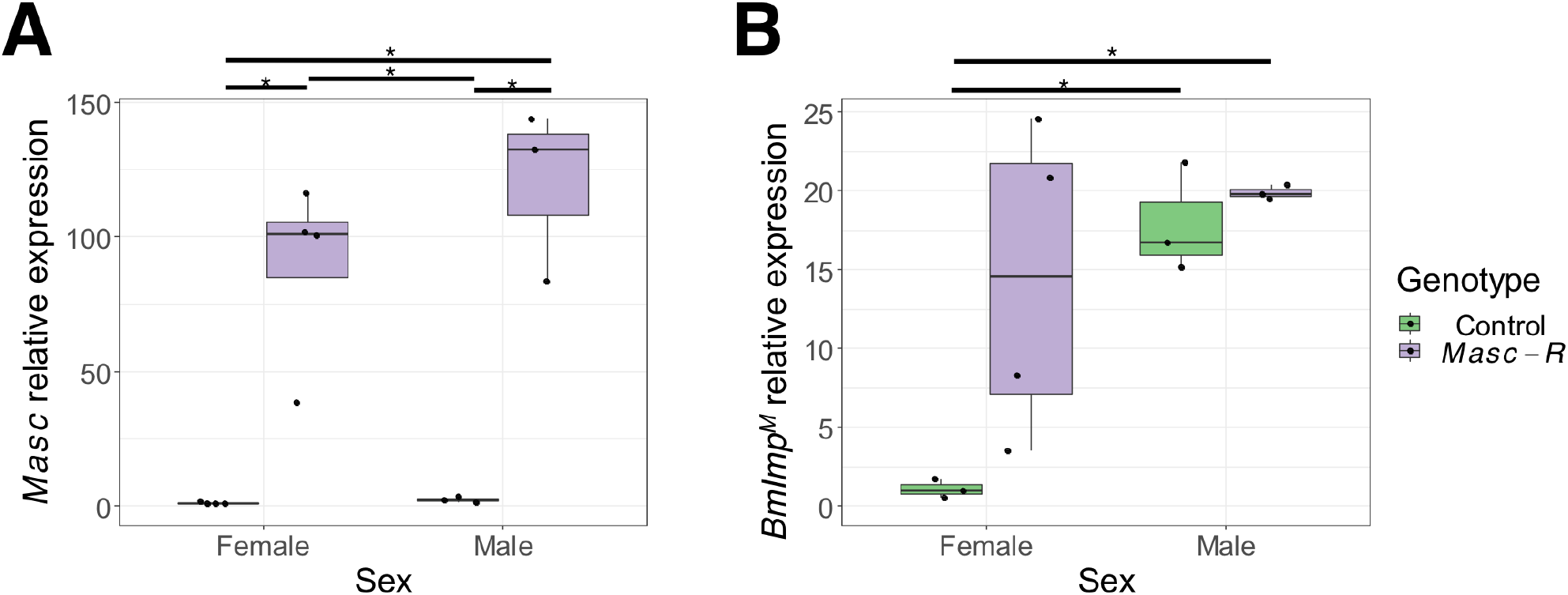
*Masc-R* overexpression-induced masculinization in females but not in males. Quantification of (A) *Masc* and (B) *BmImp^M^* mRNA. The relative mRNA levels (control female = 1) were normalized to that of *ribosomal protein 49* (*rp49*) The five points of boxplots represent the maximum, 75th percentile, 50th percentile (median), 25th percentile, and minimum value from top to bottom. The number of replicates is shown in Table S2. Asterisks indicate a significant difference in Tukey’s honestly significant difference test (p < 0.05).

Next, RNA-seq was performed using total RNA extracted from *Masc-R*-overexpressing or control neonate larvae. In *B. mori* ovary-derived cultured cells, transfection of *Masc-R* cDNA results in a growth inhibition [11]. It is also suggested that the growth inhibition induced by *Masc-R* is probably not associated with apoptosis [11]. GO enrichment analysis revealed that apoptosis-related genes were not significantly differentially expressed between wild-type and *Masc-R*-overexpressing females (Fig. S1). MA plots and boxplots of the mapping results showed that *Masc-R* overexpression decreased the expression of Z chromosome-linked genes in females but not in males (Fig. 2; Tables S3 and S4). Also, Masc-regulated genes were uniformly dispersed throughout the Z chromosome (Fig. 3). In contrast, *Masc-R* overexpression did not greatly affect gene expression on chromosome 9, where *Masc-R* transgene was inserted, in either females or males (Figs. 2 and 3). In addition, 19 DEGs are commonly identified between *Masc-R*-overexpressing and control individuals in both sexes (Table 1). To validate the RNA-seq results, the relative gene expression of four representative Z-linked genes and two chromosome 9-linked (*Masc-R* neighboring) [12] genes was measured by RT-qPCR. The expression levels of all examined Z-linked genes were about twofold decreased in *Masc-R*-overexpressing females than in control females, whereas they were comparable between *Masc-R*-overexpressing and control males (Figs. S2A-D). The expression levels of *Masc-R* neighboring genes were comparable between control and *Masc-R*-overexpressing neonate larvae, although there was a sex bias (Figs. S2E and F).

**Fig. 2.**
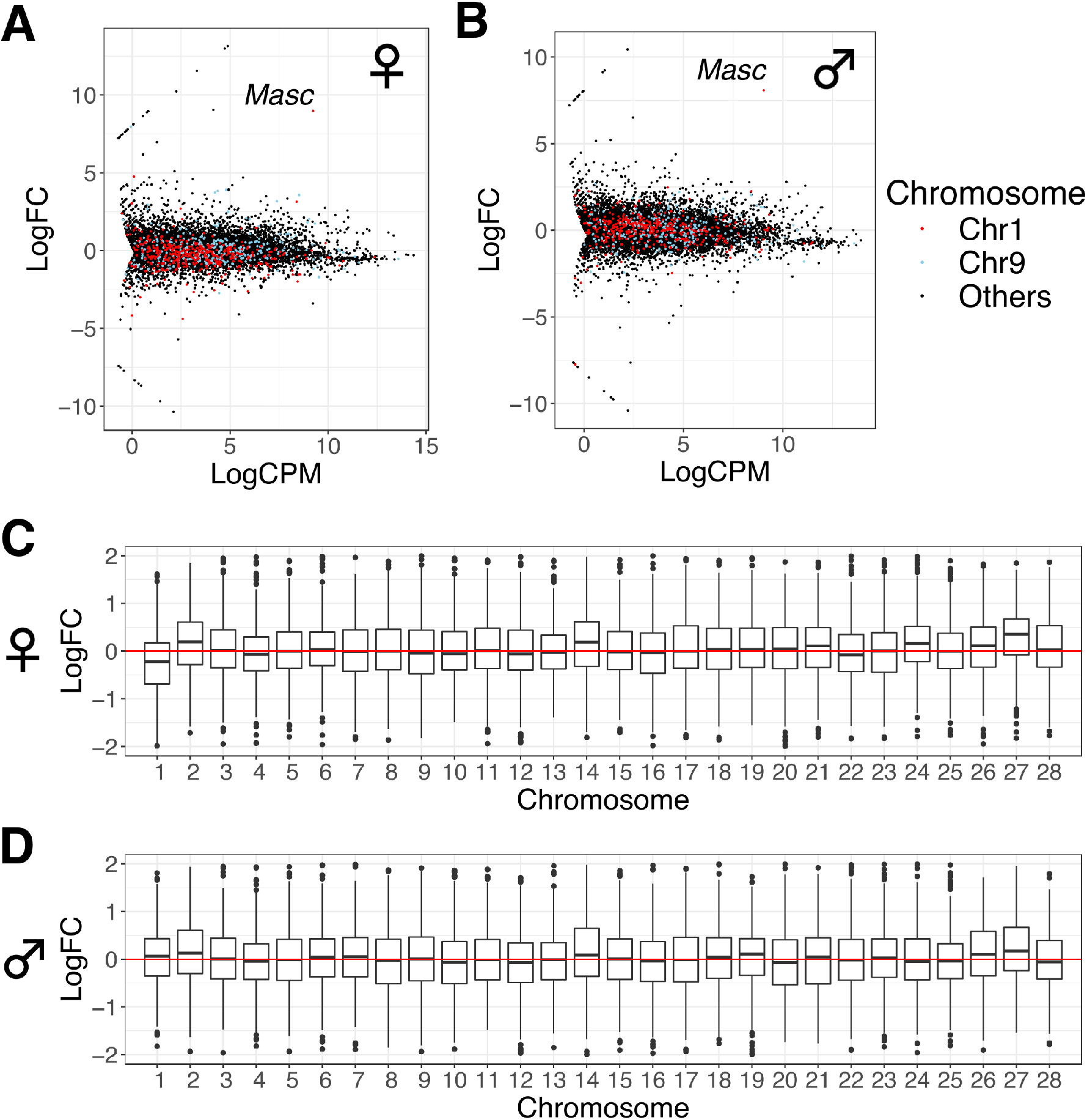
*Masc-R* overexpression-induced dosage compensation in females but not in males. (A and B) MA plots (A) between *Masc-R*-overexpressing and control females and (B) between *Masc-R*-overexpressing and control males. Z-linked genes were colored red, and chromosome 9-linked genes were colored blue. (C and D) Chromosomal distribution of differentially expressed transcripts (C) between *Masc-R*-overexpressing and control females and (D) between *Masc-R*-overexpressing and control males. logCPM: average log_2_ CPM; logFC: log_2_-fold change in expression ratios of genes in *Masc-R*-overexpressing individuals compared with control individuals.

**Fig. 3.**
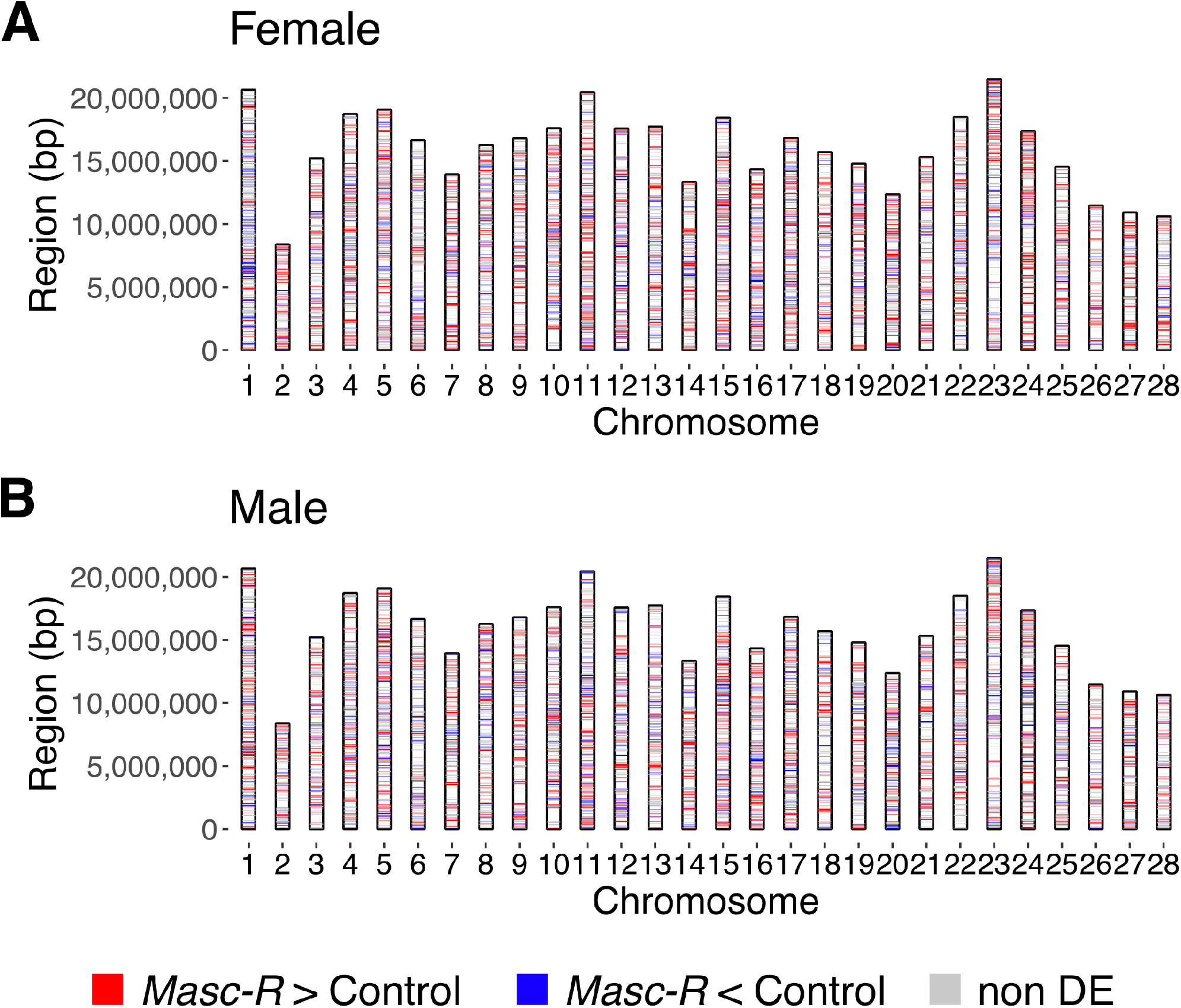
Genomic loci of Masc-regulated genes in (A) females and (B) males. logFC > 0.3 are colored red, logFC < −0.3 are colored blue, and −0.3 ≤ logFC ≤ 0.3 are colored gray.

**Table 1.**
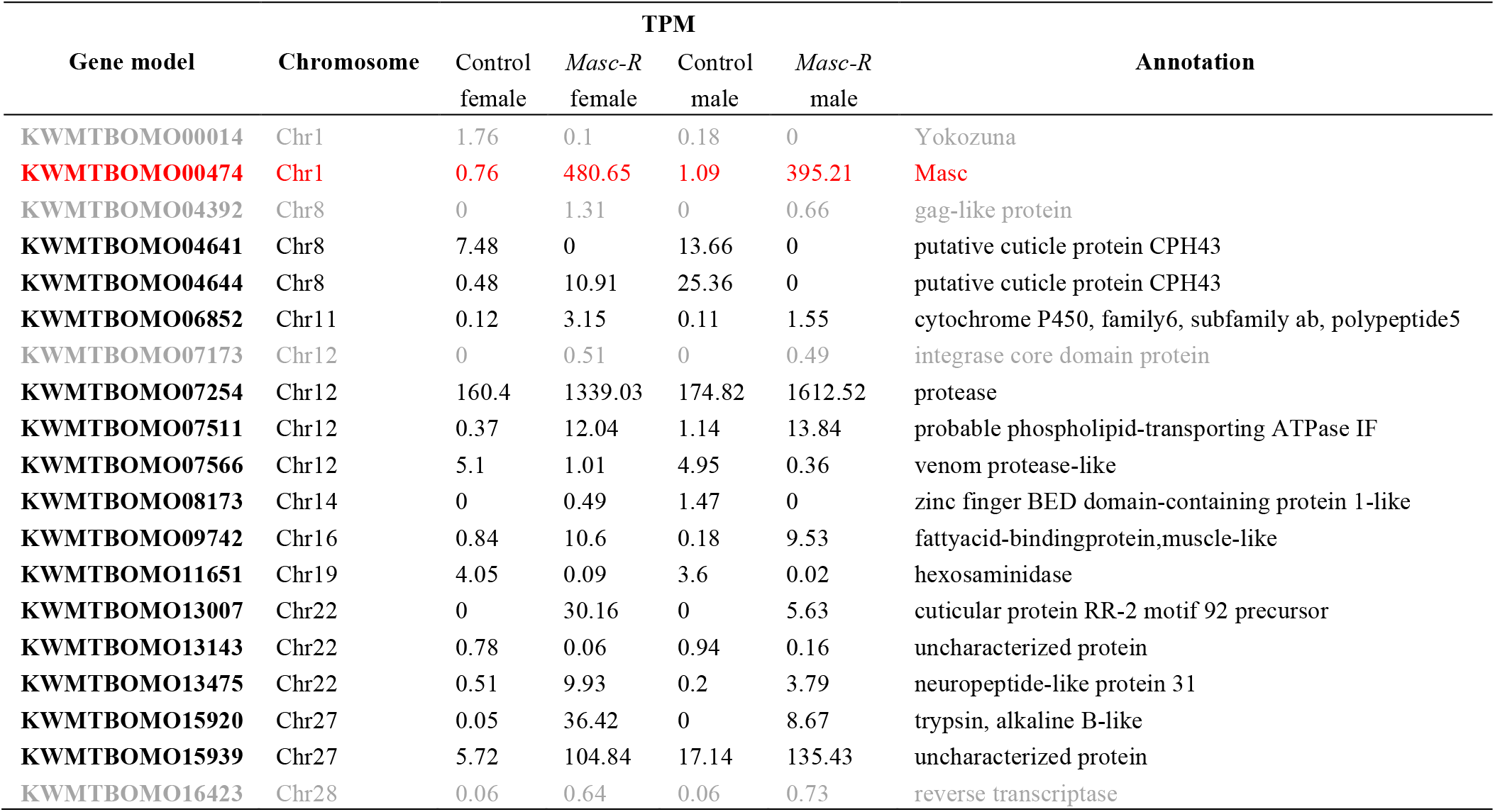
DEGs that were common between *Masc-R*-overexpressing and control individuals in both sexes. *Masc* is shown in red letters. Putative transposable elements are shown in gray letters. TPM: transcripts per million.

| Gene model | Chromosome | TPM |  |  |  | Annotation |
| --- | --- | --- | --- | --- | --- | --- |
|  |  | Control female | *Masc-R* female | Control male | *Masc-R* male |  |
| KWMTBOMO00014 | Chr1 | 1.76 | 0.1 | 0.18 | 0 | Yokozuna |
| **KWMTBOMO00474** | **Chr1** | **0.76** | **480.65** | **1.09** | **395.21** | **Masc** |
| KWMTBOMO04392 | Chr8 | 0 | 1.31 | 0 | 0.66 | gag-like protein |
| KWMTBOMO04641 | Chr8 | 7.48 | 0 | 13.66 | 0 | putative cuticle protein CPH43 |
| KWMTBOMO04644 | Chr8 | 0.48 | 10.91 | 25.36 | 0 | putative cuticle protein CPH43 |
| KWMTBOMO06852 | Chr11 | 0.12 | 3.15 | 0.11 | 1.55 | cytochrome P450, family6, subfamily ab, polypeptide5 |
| KWMTBOMO07173 | Chr12 | 0 | 0.51 | 0 | 0.49 | integrase core domain protein |
| KWMTBOMO07254 | Chr12 | 160.4 | 1339.03 | 174.82 | 1612.52 | protease |
| KWMTBOMO07511 | Chr12 | 0.37 | 12.04 | 1.14 | 13.84 | probable phospholipid-transporting ATPase IF |
| KWMTBOMO07566 | Chr12 | 5.1 | 1.01 | 4.95 | 0.36 | venom protease-like |
| KWMTBOMO08173 | Chr14 | 0 | 0.49 | 1.47 | 0 | zinc finger BED domain-containing protein 1-like |
| KWMTBOMO09742 | Chr16 | 0.84 | 10.6 | 0.18 | 9.53 | fattyacid-bindingprotein,muscle-like |
| KWMTBOMO11651 | Chr19 | 4.05 | 0.09 | 3.6 | 0.02 | hexosaminidase |
| KWMTBOMO13007 | Chr22 | 0 | 30.16 | 0 | 5.63 | cuticular protein RR-2 motif 92 precursor |
| KWMTBOMO13143 | Chr22 | 0.78 | 0.06 | 0.94 | 0.16 | uncharacterized protein |
| KWMTBOMO13475 | Chr22 | 0.51 | 9.93 | 0.2 | 3.79 | neuropeptide-like protein 31 |
| KWMTBOMO15920 | Chr27 | 0.05 | 36.42 | 0 | 8.67 | trypsin, alkaline B-like |
| KWMTBOMO15939 | Chr27 | 5.72 | 104.84 | 17.14 | 135.43 | uncharacterized protein |
| KWMTBOMO16423 | Chr28 | 0.06 | 0.64 | 0.06 | 0.73 | reverse transcriptase |

## Discussion

This study showed that the expression of Z-linked genes is globally suppressed in *Masc-R*-overexpressing females (Figs. 2 and 3). This finding was consistent with previous studies demonstrating the global hyperactivation of Z-linked genes in *Masc* knocked down males [8,10]. Results also indicated that *Masc-R* overexpression is sufficient for the induction of dosage compensation in females, and elements necessary for dosage compensation are likely to be present in females except for Masc. In contrast, *Masc-R* overexpression did not greatly affect gene expression on chromosome 9, where *Masc-R* transgene is inserted (Figs. 2, 3, and S2). This indicated that the location where *Masc* is transcribed is not associated with dosage compensation and that dosage compensation factors, which are not identified in *B. mori*, only repress the expression of Z-linked genes. Thus, dosage compensation factors recognize the same features of the Z chromosome in both sexes.

In *Masc-R*-overexpressing female neonates, 65.7% of the Z-linked genes were not downregulated (logFC > −0.5 in Table S3). One possible explanation for this insufficiency of *Masc*-induced dosage compensation is a limitation of the *BmA3* promoter, which was used to induce *GAL4* expression. Although the *BmA3* promoter induces widespread gene expression, it is not universal and exhibits little to no activity in several larval tissues and embryos [22,23] and is susceptible to positional effects [14,23]. Sakai et al. [12] reported that BmA3-GAL4*-*induced expression of *Masc-R* resulted in the expression of both female and male forms of *doublesex* transcripts (*dsx^F^* and *dsx^M^*) [24,25] in neonate larvae, supporting the insufficiency of *Masc*-induced masculinization driven by the *BmA3* promoter. Using other ubiquitous promoters, such as hsp90^P2.9k^ [26], dosage compensation in *Masc-R*-overexpressing females may be fully established. Another possibility is the accumulation of male-biased genes on the Z chromosome in species with a WZ sex-determination system [27]. Previous studies in several lepidopteran species reported the male-biased expression of the Z chromosome [28–30]. These male-biased genes are presumed to be hyperactivated in the presence of Masc and behave as if they escaped from dosage compensation. The Z-linked male-specific transcript, *BmImp^M^*, was not downregulated but upregulated in *Masc-R*-overexpressing females (Fig. 1B).

Sakai et al. [12] reported that *B. mori* females overexpressing *Masc-R* under the control of the *BmA3* promoter are dead during the larval stage, but the cause of this lethality remains unclear. In *B. mori* ovary-derived cultured cells, transfection of *Masc-R* cDNA resulted in a growth inhibition [11]. Because caspase activity is not upregulated in *Masc-R*-transfected cells, cell growth inhibition induced by *Masc-R* is probably not associated with apoptosis [11]. GO enrichment analysis was performed in *Masc-R*-overexpressing females to confirm the previous observation *in vitro*: apoptosis-related genes were not significantly differentially expressed between control and *Masc-R*-overexpressing females (Fig. S1). This suggested that lethality is not caused by apoptosis-mediated cell death. In *D. melanogaster*, *C. elegans*, and the mouse *Mus musculus*, abnormal expression of X-linked genes caused by a failure of dosage compensation resulted in sex-specific lethality during the developmental stage [31–33]. In *B. mori, Masc* knocked down males were lethal at the embryonic stage [10]. In addition, in *D. melanogaster*, forced expression of *msl2* that encoded a subunit of dosage compensation complex caused female-specific lethality [1,34,35]. This study observed an abnormal expression of Z-linked genes in *Masc-R*-overexpressing females (Figs. 2 and 3). Thus, although there was no direct evidence, the female-specific lethality observed in *Masc-R*-overexpressing larvae was probably due to the abnormal expression of Z-linked genes in females.

*Masc-R*-overexpressing males did not show any abnormalities in growth, development, and sexual dimorphisms [12]. Although *Masc* expression was 50-fold higher in *Masc-R*-overexpressing males than in control males (Fig. 1A), the expression of Z-linked genes (Figs. 2 and 3; Table S4) and *BmImp^M^* was comparable to that of control males (Fig. 1B). These results indicated that excessive *Masc* expression strengthens neither dosage compensation nor maleness in males. In contrast, 94 DEGs between *Masc-R*-overexpressing and control males were identified (Table S4); of these, 19 DEGs were common in both sexes (Table 1). One of the DEGs was *Masc*, four DEGs encoded putative transposable elements, and the other 14 DEGs encoded functional proteins. In addition, although the reason is unclear, *Masc-R* overexpression increased gene expression on chromosome 27 in both sexes (Fig. 2; Tables S3 and S4). These genes were probably directly or indirectly regulated by Masc and may have functions other than sex-determination and dosage compensation. *Sex-determining region on Y chromosome* (*SRY*) was overexpressed in ~84% of male hepatocellular carcinoma patients, and *Sry* overexpression in *M. musculus* resulted in hepatocarcinogenesis [36]. In *D. melanogaster*, the master-feminizing gene *Sxl* also functioned in germ cell meiosis and differentiation [37]. According to the *B. mori* genome database (SilkDB 3.0 [38]) and a previous study [39], *Masc* is expressed in both somatic and germ tissues. Additionally, Masc is involved in developing the female external genitalia [40], suggesting that Masc has some unknown functions in both sexes. By observing in detail the phenotypes of *Masc-R*-overexpressing individuals and investigating the functions of genes expressed under the control of Masc (Tables 1, S3, and S4), additional functions of Masc may be revealed.

## Acknowledgements

We thank the Institute for Sustainable Agro-ecosystem Services, The University of Tokyo, for facilitating the mulberry cultivation and the Biotron Facility at the University of Tokyo for rearing the silkworms. We thank Drs. Hiroyuki Hikida and Keisuke Shoji for their useful comments. We are grateful to Wakako Saito and Natsuki Nakashima for technical assistance for silkworm maintenance. This work was supported by grants from MEXT KAKENHI 17H06431 to SK and TK.

## Author contributions

TK and SK designed the study. TK prepared insects. KT and TK performed the molecular experiments. KT analyzed the data. KT wrote the draft manuscript and TK revised it with the inputs of SK.

## Data accessibility

RNA-seq data was submitted to DDBJ under the accession number XXXX.

## Supplementary Material

**Table S1.** Primer lists used in this study.

**Table S2.** Number of individuals used in RT-qPCR and RNA-seq analysis.

**Table S3.** DE analysis between *Masc-R*-overexpressing and control females. TPM: transcripts per million.

**Table S4.** DE analysis between *Masc-R*-overexpressing and control males. The abbreviation is described in Table S3.

**Fig. S1.** Significantly enriched GO terms of DEGs between *Masc-R*-overexpressing and control females. Numbers on the bar graph indicate DEGs (FDR < 0.05) out of all genes classified to the corresponding GO term. BP: biological process, CC: cellular component, MF: molecular function.

**Fig. S2.** Effects of *Masc-R* overexpression on (A–D) Z-linked genes and (E and F) *Masc-R* neighboring genes. Quantification of (A) *KWMTBOMO00646*, (B) *KWMTBOMO00338*, (C) *KWMTBOMO00238*, (D) *KWMTBOMO00345*, (E) *KWMTBOMO05086*, and (F) *KWMTBOMO05084* mRNA. The relative mRNA levels (control female = 1) were normalized to that of *rp49*. The explanation of boxplots is described in Fig. 1. The number of replicates is shown in Table S2. Asterisks indicate a significant difference in Tukey’s honestly significant difference test (p < 0.05).

