## Supplemental figures for "Transcriptome analysis in the silkworm *Bombyx mori* overexpressing piRNA-resistant *Masculinizer* gene"

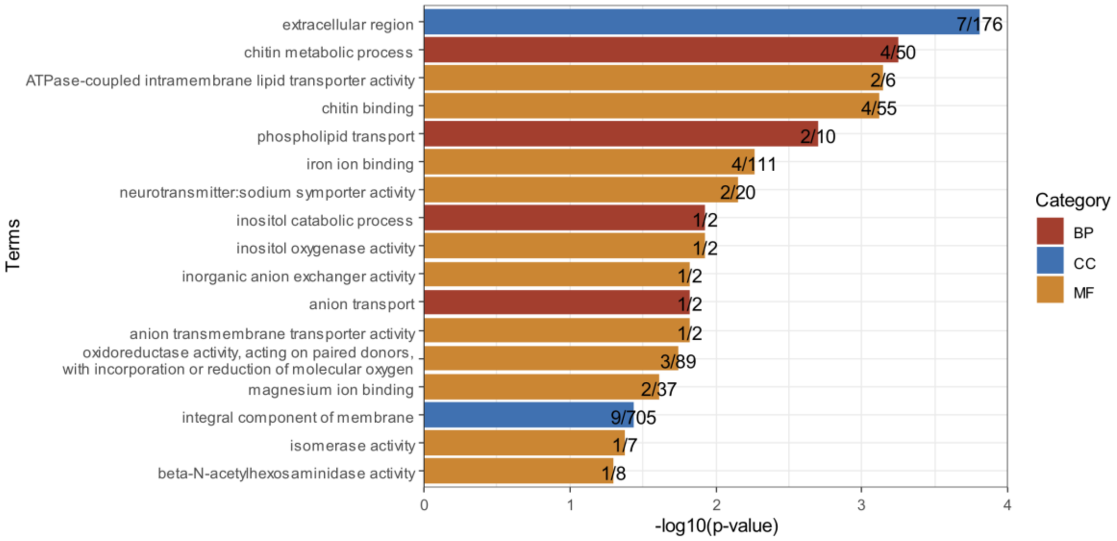


**Fig. S1.** Significantly enriched GO terms of DEGs between *Masc-R*-overexpressing and control females. Numbers on the bar graph indicate DEGs (FDR < 0.05) out of all genes classified to the corresponding GO term. BP: biological process, CC: cellular component, MF: molecular function.

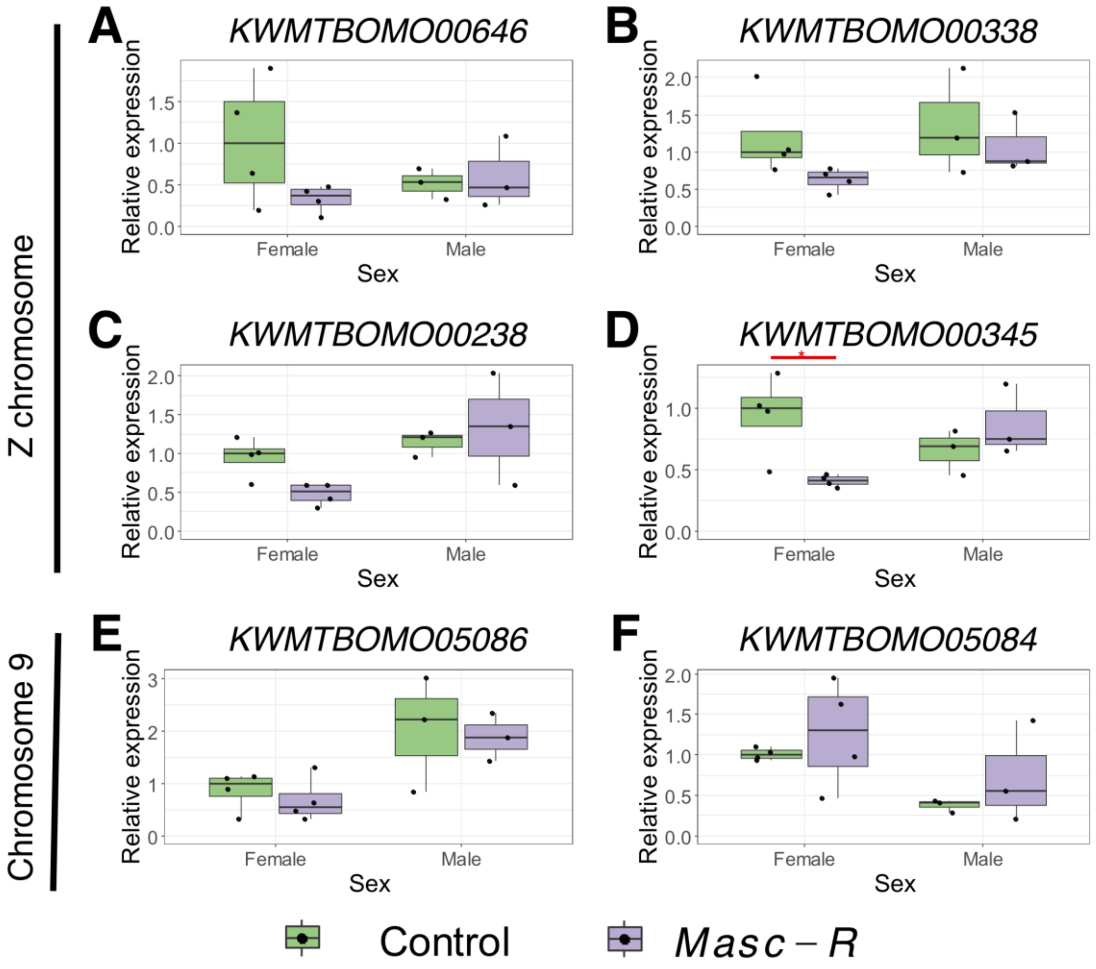


**Fig. S2.** Effects of *Masc-R* overexpression on (A–D) Z-linked genes and (E and F) *Masc-R* neighboring genes. Quantification of (A) *KWMTBOMO00646*, (B) *KWMTBOMO00338*, (C) *KWMTBOMO00238*, (D) *KWMTBOMO00345*, (E) *KWMTBOMO05086*, and (F) *KWMTBOMO05084* mRNA. The relative mRNA levels (control female = 1) were normalized to that of *rp49*. The explanation of boxplots is described in Fig. 1. The number of replicates is shown in Table S2. Asterisks indicate a significant difference in Tukey's honestly significant difference test ($p < 0.05$).
